# A functional circadian clock regulates composition and daily bacterial load of the gut microbiome in *Drosophila melanogaster*

**DOI:** 10.1101/2024.03.15.585158

**Authors:** Matteo Battistolli, Irene Varponi, Ottavia Romoli, Federica Sandrelli

## Abstract

While *Drosophila melanogaster* serves as a crucial model for investigating both the circadian clock and gut microbiome, our understanding of their relationship in this organism is still limited. Recent analyses suggested that the *Drosophila* gut microbiome modulates the host circadian tran-scriptome to minimize rapid oscillations in response to changing environments. To delve deeper into the potential relationship between the gut microbiota and circadian clock in *Drosophila*, we examined the composition and abundance of the gut microbiota in wild-type and arrhythmic *per*^01^ flies, under 12 h:12 h light: dark (12:12 LD) and constant darkness (DD) conditions. The gut microbiota of wild-type and *per*^01^ flies showed differences in composition, suggesting that the *D. melanogaster* circadian clock has a role in shaping the gut microbiome. In 12:12 LD and DD conditions, *per*^01^ mutants showed significant daily variations in gut bacterial quantity, unlike wild-type flies. This suggests that the circadian clock in *D. melanogaster* plays a role in maintaining daily stability in gut microbiome load. Finally, some gut bacteria exhibited significant 24 h fluctuations in their relative abundance, which appeared independent from the fly circadian clock, suggesting that certain gut commensal bacteria in *Drosophila* may possess a host-independent circadian clock.

## 1. Introduction

Enteric bacteria deeply impact various host physiological processes, including metabolism, energy homeostasis, immune response, as well as several neurological processes, establishing a complex bidirectional host-microbe relationship ^1, 2^. Recognized as a “virtual organ within an organ” ^3^, the gut microbiota plays a pivotal role in regulating the host’s overall health. Disruptions in gut microbiota composition, or dysbiosis, along with subsequent disturbances in microbiota metabolic activity, have been linked to numerous human pathologies, including obesity, immune-related diseases, and mood disorders ^2^. Both genetic factors of the host and environmental influences such as diet and lifestyle contribute to shaping the composition of the gut microbiome. However, the relative contributions of these factors and their potential interplay in regulating the gut microbiota remain incompletely understood ^4–6^. An increasing body of evidence indicates that the host’s circadian clock has an important role in modulating the relationship between the host and gut microbiome in both health and disease states ^7–9^.

Circadian clocks are genetically controlled timekeeping mechanisms which enable organisms to coordinate their physiological and behavioral activities with the daily environmental variations caused by the Earth’s rotation. These endogenous clocks can measure time in a 24 h temporal domain, in the absence of any environmental stimulus (free running conditions); additionally, they can be synchronized (entrained) by external cues, such as day-night light variations, allowing organisms to fine-tune their physiology and behavior in phase with the 24 h day ^10,11^. At a molecular level, circadian clocks are based on auto-regulatory and interlocked transcriptional-translational feedback loops, in which positive elements promote the production of their inhibitors. These cycling molecular oscillations control the transcriptional program of cells and tissues in turn generating rhythmic activities at an organismal level ^12^.

Several studies indicate that a robust interconnection between the gut microbiota and the host’s circadian clock exists. For example, both human and mouse models exhibit daily rhythmicity in gut microbiota composition and metabolite levels, which can be perturbed by genetic or environmental disruptions of the circadian clock ^13–17^. On the other hand, the gut microbiota was shown to play a fundamental role in regulating host’s circadian transcriptional profiles in the gut and liver, thereby modulating the circadian regulation of host metabolism ^14,18–20^. A recent study in mice demonstrated that the gut microbiota promotes daily innate immunity oscillations that correlate with feeding rhythms, thus anticipating possible oral exposure to pathogens like *Salmonella* Typhimurium^21^. Additionally, different studies suggest a link between disruptions in the circadian clock and dysbiosis detected in various human pathologies, including obesity, diabetes, cardiovascular diseases, neurodegenerative disorders, and sensitivity to infections ^7^.

*Drosophila melanogaster* has been fundamental for deciphering the circadian system at the molecular/cellular and organismal levels ^12^. With its relatively simple gut microbiota, comprising a limited number of species, the fruit fly serves as an excellent model organism for investigating how the microbiota affects multiple aspects of host physiology, including development, lifespan, immune response, and some behavioral phenotypes ^22–28^.

Despite *D. melanogaster* significance in studying both the gut microbiota and circadian clock, scant data exist on the potential interplay between these two aspects. Only one study has been published thus far on this subject ^29^. Through an extensive transcriptomic analysis of guts from flies reared under sterile and normal conditions and subjected to different feeding regimes, this work revealed that the gut microbiome has a role in stabilizing daily gut transcriptome oscillations, likely favoring circadian synchrony among different organs at an organismal level ^29^. Additionally, Zhang and colleagues examined daily variations in the with-in structure of microbiota in wild type and *per* ^01^ flies, which carry a null mutation in the cardinal clock gene *period* (*per*). These flies were either fed *ad libitum* or subject to a timed feeding (TF) paradigm, known to promote daily rhythmicity in mammals ^13,29^. Notably, significant cycling was exclusively observed in the microbiome of *per* ^01^ flies under TF regimes and was restricted to only two bacterial taxa, suggesting that *Drosophila* gut microbiome composition does not exhibit daily cycling, unlike mammals ^29^. However, since these analyses focused on microbiome derived from fecal samples of flies reared under 12 h: 12 h light: dark (12:12 LD) conditions, additional data are required to elucidate the relationship between the circadian clock and gut microbiota in flies.

Here we analyzed the composition and daily abundance of microbiota derived from dissected guts of wild-type flies and isogenic *per*^01^ arrhythmic mutants, reared in parallel in 12:12 LD and constant darkness (DD) conditions, and fed *ad libitum* on the same diet. Wild-type and *per*^01^ flies showed differences in gut microbiota composition, suggesting that as in mammals the *D. melanogaster* circadian clock has a role in shaping gut microbiome. Additionally, our data indicate that a functional clock inhibits daily oscillations in the gut bacterial total abundance in both 12: 12 LD and DD regimes, suggesting that the circadian clock is important for guarantying a daily stability of the gut microbiome levels in *D. melanogaster*. Nevertheless, few components of the gut microbiota of both wild-type and arrhythmic *per*^01^ flies showed significant daily fluctuations in their relative abundance, possibly suggesting that some gut commensal bacteria may have a host-independent circadian clock in *D. melanogaster*.

## 2. Results

### 2.1 In *Drosophila melanogaster* a functional circadian clock stabilizes gut bacterial levels throughout the day

In this study we analyzed gut microbiome composition and abundance in wild-type (Canton-S) and circadian clock *per*^01^ mutant flies, under both 12:12 LD and DD conditions. To limit possible confounding effects due to variations in genetic and environmental factors, we used arrhythmic *per*^01^ flies belonging to a cantonized strain. These flies carry a null mutation at the level of the *per* gene and have the same genetic background as the *per*^+^ wild-type strain (Supplementary Fig. S1a,b) ^30^). Both *Drosophila* strains were free from any *Wolbachia* spp. (Supplementary Fig. S1c), a bacterial endosymbiont whose presence could affect the gut microbiota sampling depth and quantification ^31^. Furthermore, all experiments investigating the microbiome were performed on wild-type and *per*^01^ males, reared in parallel under identical environmental conditions and provided with the same food source.

In 12:12 LD or at the third day of DD, guts derived from four-seven day-old Canton-S and *per*^01^ males were collected at Zeitgeber Times (ZTs) or Circadian Times (CTs) 0.5, 6, 12.5, and 18 (with ZTs 0 and 12 respectively corresponding to light-on and -off, in 12:12 LD; and CTs 0 and 12 corresponding to the beginning and end of the subjective day, in DD conditions). Gut samples were analyzed in parallel for estimation of total bacterial abundance and *16S* rDNA sequencing (Fig. 1a). Quantification of bacterial abundance via quantitative PCR (qPCR) did not reveal any significant daily oscillation in the total microbiota load of wild-type flies, in either 12:12 LD or DD conditions [Fig. 1b,c; P> 0.05, not significant (ns), for both 12:12 LD and DD regimes]. Conversely, *per*^01^ flies showed significant daily variations in bacterial abundances, with low microbiota loads at ZT/CT 18, under both 12:12 LD and DD conditions (Fig. 1d,e; P< 0.05 in both lighting conditions).

**Figure 1.**
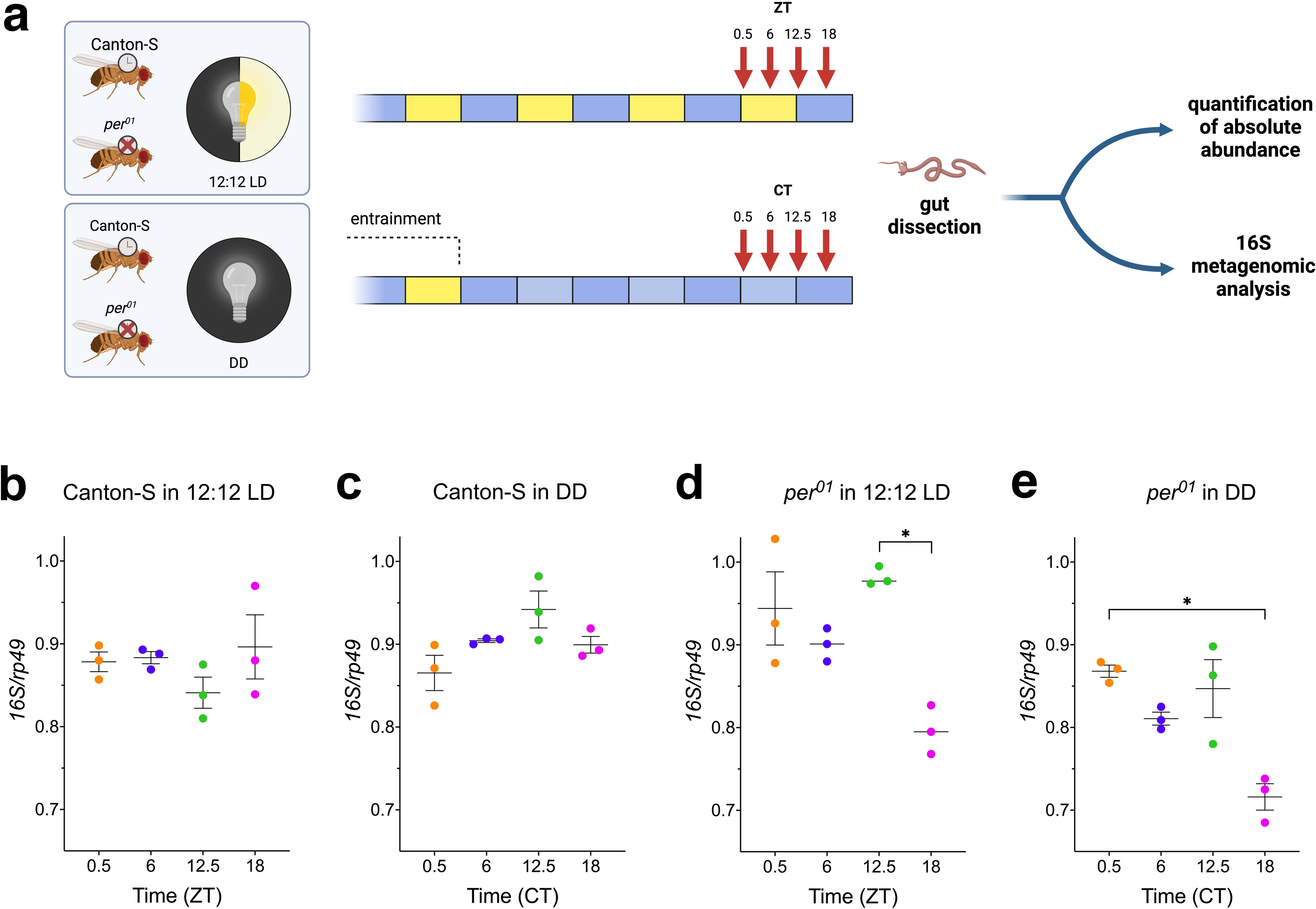
Experimental design and total gut bacteria abundances in wild-type and *per^01^*flies under 12:12 LD and DD conditions. (**a**) Experimental setup for the characterization of gut microbiota in wild-type (Canton-S) and *per^01^*males under 12:12 LD and DD regimes. In 12:12 LD conditions, flies were collected at 5-7 days post-eclosion. For the collection in DD, newly eclosed flies were entrained at least three days under 12:12 LD regime and transferred to DD two days before sampling. Red arrows indicate the time points at which sampling occurred (ZTs/CTs 0.5, 6, 12.5, and 18). For each genotype and lighting condition, 20 to 60 guts per time point (in three replicates) were dissected. Gut samples were processed in parallel for the evaluation of total gut bacterial abundance and *16S* rDNA sequencing (details on collected samples in Supplementary Table S1). This illustration was created with BioRender.com. (**b-e**) Daily variation of total microbial abundances (mean ± SEM, N= 3 per time point) in Canton-S and *per^01^* guts under 12:12 LD and DD conditions. Bacterial DNA was quantified via qPCR on genomic DNA obtained from Canton-S and *per^01^* guts. *Drosophila rp49* was used to normalize the amount of detected *16S* rDNA. Non-parametric one-way ANOVA (Kruskal-Wallis test) showed that Canton-S flies maintained stable gut microbiota levels throughout the day under both (**b**) 12:12 LD (P = 0.31, not significant, ns) and (**c**) DD (P = 0.06, ns), while *per^01^* flies exhibited daily fluctuations in gut bacterial content in both (**d**) 12:12 LD (P = 0.017) and (**e**) DD conditions (P = 0.020). * indicates: P = 0.039 (**d**) and P = 0.028 (**e**) in Dunn’s *post hoc* test.

### 2.2 The *Drosophila* circadian clock shapes gut microbiota composition

We next examined wild-type and *per*^01^ gut microbial compositions in 12:12 LD and DD regimes, via *16S* profiling. A total number of 1919 Amplicon Sequence Variants (ASVs) passed the filtering step and were used to define the rarefied dataset (Supplementary Fig. S2 and Supplementary Table S1). Canton-S and *per*^01^ guts shared a total of 46 ASVs, with only 11 ASVs common to the two host genotypes under both 12:12 LD and DD conditions. Interestingly, 1073 and 468 ASVs were exclusively found in wild-type and mutant flies, respectively (Fig. 2a).

**Figure 2:**
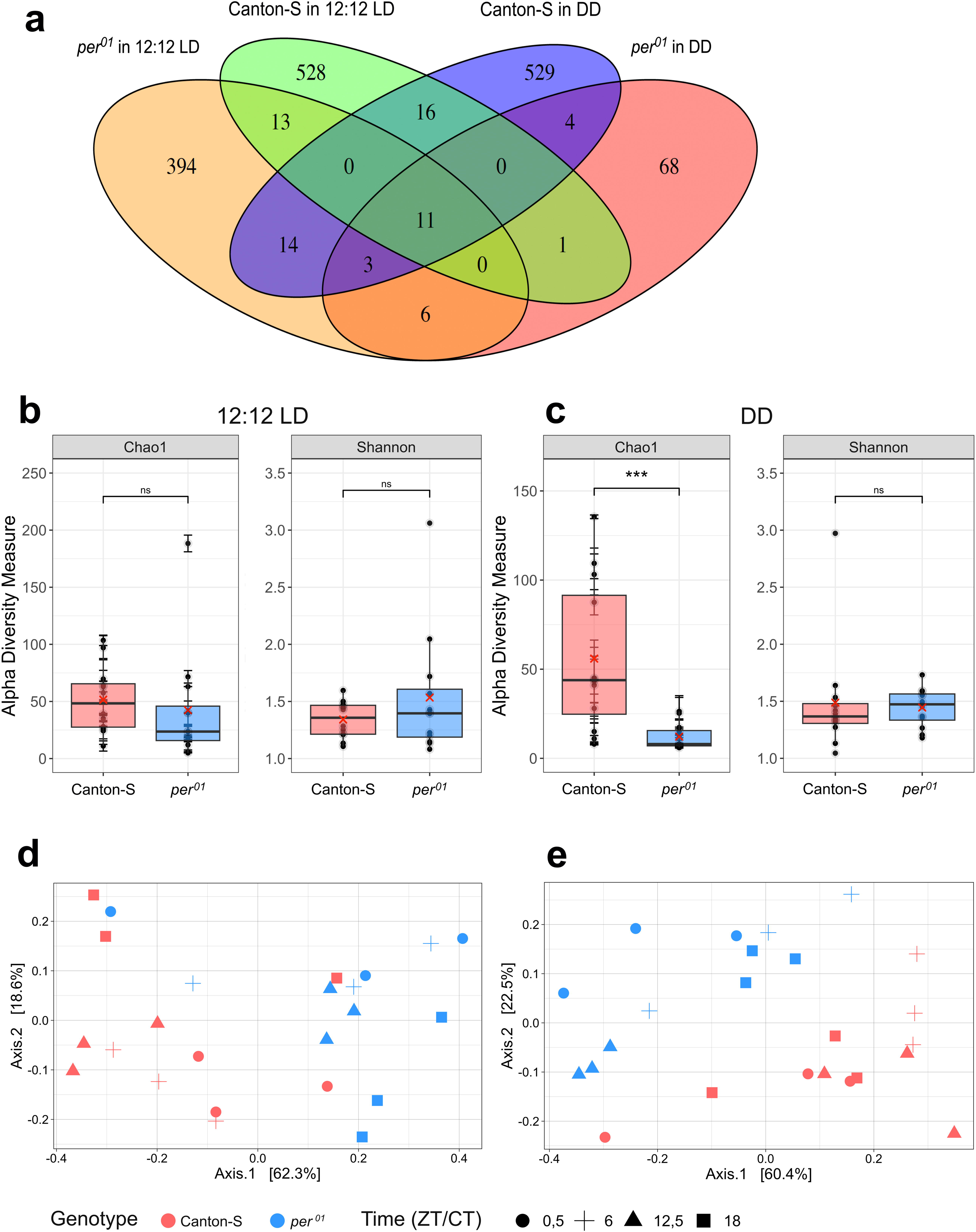
Comparison of gut microbiota in wild-type and *per^01^* flies under 12:12 LD and DD conditions. (**a**) Venn diagram illustrating the number of ASVs unique or shared between the different genotypes (Canton-S and *per^01^*) and lighting conditions (12:12 LD and DD). (**b,c**) Comparisons of Shannon and Chao1 alpha diversity indices between Canton-S and *per^01^* microbiota under (**b**) 12:12 LD or (**c**) DD conditions. In each box plot, mean and median values are shown by red Xs and horizontal black lines, respectively; each dot represents a biological replicate. In (**b**), no significant differences in Chao1 and Shannon indices were detected between genotypes (Chao1: P = 0.133, ns; Shannon: P = 0.862, ns; Kruskal-Wallis test). In (**c**), Chao1 (P = 0.0009) but not the Shannon index (P = 0.419, ns) resulted significantly different between the two genotypes. (**d, e**) Principal Coordinates Analysis (PCoA) of Bray-Curtis distances of Canton-S (red) and *per^01^*(blue) microbiota under (**d**) 12:12 LD and (**e**) DD regimes. Different symbols indicate different time points: circle = ZT/CT 0.5, cross = ZT/CT 6, triangle = ZT/CT 12.5, square = ZT/CT 18. PERMANOVA analyses showed significant differences in gut microbiota composition of Canton-S and *per^01^*, in (**d**) 12:12 LD (P = 0.001), and (**e**) DD (P = 0.001) conditions.

We used alpha diversity metrics to evaluate whether wild-type and *per*^01^ flies showed some differences in the with-in sample structure of their gut microbiota in 12: 12 LD and DD conditions. In 12:12 LD regimes, wild-type and *per*^01^ gut microbiota showed similar total species richness and evenness (Fig. 2b; P > 0.05, ns for Chao1 and Shannon indices). In DD conditions, some differences between the microbiota of the two host genotypes were detected since the total species richness resulted significantly lower in *per*^01^ flies compared to wild-type individuals (Fig. 2c; Chao1 index: P = 0.0009), although the species richness and evenness, evaluated together by the Shannon index were similar between the two genotypes (Fig. 2c; P > 0.05, ns). Subsequently, we compared the gut microbiota composition in wild-type and *per*^01^ flies, performing a beta diversity analysis. We found significant dissimilarities in microbial composition between the two host genotypes under both 12:12 LD and DD conditions (Fig. 2d,e; P = 0.001 in both 12:12 LD and DD regimes).

To understand the reason behind these variations, we first examined the distribution of the microbiota most prevalent bacterial families in the two host genotypes. When considering the entire set of wild-type and *per*^01^ gut samples, the most abundant families were *Lactobacillaceae* (min 35.54 %, max 93.58 %) and *Acetobacteraceae* (min 5.5 %, max 57.34 %). Together these two families represented ∼ 97 % of the average relative abundance (min 68.73%, max 100%) and displayed comparable levels in Canton-S and *per*^01^ flies (Fig. 3a,b; P > 0.05, ns, for both comparisons). Subsequently, we focused on the lowest taxonomic level, selecting the most abundant ASVs that collectively represented ∼ 93% of the whole microbial diversity among all samples. Out of these most prevalent ASVs, three corresponded to *Lactiplantibacillus plantarum*. When we analyzed the *16S* rDNA sequence of these three ASVs, we found that they differed for a single nucleotide modification in distinct positions of the analyzed *16S* rDNA portion (Supplementary Fig. S3). Since *L. plantarum* was reported to contain various *16S* rDNA copies ^32,33^, the three different *16S* DNA sequences might result from distinct ribosomal RNA operons within the same bacterial strain. Therefore, we grouped the three *L. plantarum* ASVs, named A, B, and C, in a single *L. plantarum* ASV cluster. Thus, among all the gut samples, the most abundant ASVs were the *Lactiplantibacillus plantarum* cluster, showing the highest prevalence (41.68 %), followed by the ASVs assigned to *Lactobacillus fructivorans* (26.86 %), *Acetobacter pasteurianus* (18.32 %), *Acetobacter malorum* (9.36 %), *Morganella morganii* (0.90 %), and *Levilactobacillus brevis* (0.31 %) (Fig. 3c).

**Figure 3:**
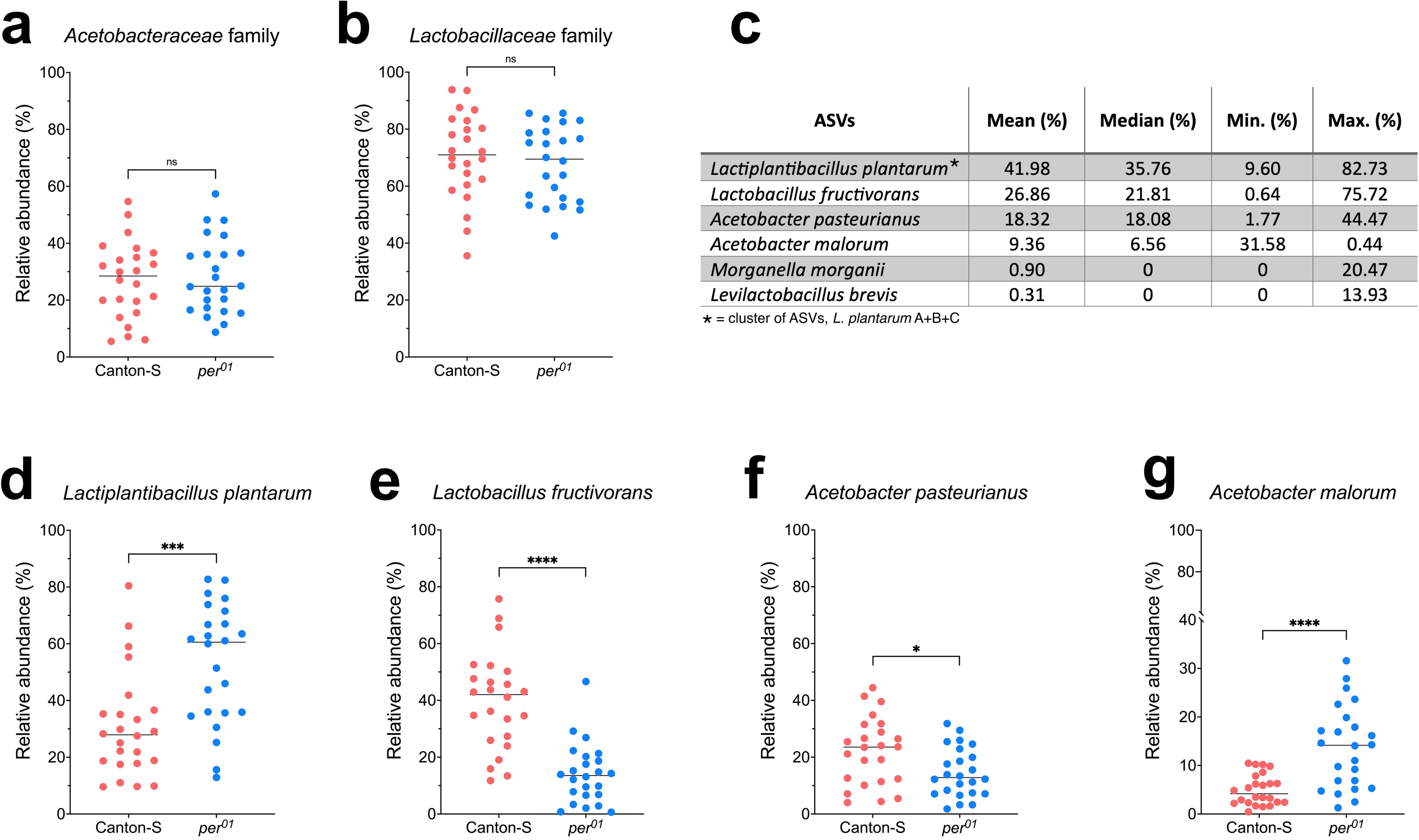
Relative abundances of the most prevalent bacterial families and ASVs in wild-type and *per^01^*microbiota. Relative abundances of (**a**) *Acetobacteraceae* and (**b**) *Lactobacillaceae* families in wild-type (Canton-S) and *per^01^* microbiota. Solid black lines represent median values. No significant differences between the two host genotypes were detected for both families [(**a**) P = 0.74, ns, and (**b**) P = 0.50, ns, in t-test]. (**c**) Mean, median, minimum, and maximum relative abundances (%) of the most prevalent ASVs among all samples. (**d-g**) Comparison of relative abundances of (**d**) *L. plantarum*, (**e**) *L. fructivorans*, (**f**) *A. pasteurianus*, and (**g**) *A. malorum* ASVs in Canton-S and *per^01^* microbiota. (**d**) *\*\*\**: P = 0.0003; (**e**) ****: P < 0.0001; (**f**) *: P = 0.017; (g) ****: P < 0.0001 in t-test.

When we compared these prevalent ASVs between the two host genotypes, two were exclusively found in *per*^01^ mutants but not in wild-type flies (*M. morganii*, *L. brevis*). The remaining ASVs (*L. plantarum* cluster, *L. fructivorans*, *A. pasteurianus*, and *A. malorum*) exhibited significant differences between wild-type and clock mutant guts (Fig. 3d-g; P< 0.05 for all comparisons). Specifically, Canton-S flies showed higher levels of *L. fructivorans* and *A. pasteurianus* (Fig. 3e,f), while *per*^01^ flies showed a greater abundance of *L. plantarum* and *A. malorum* (Fig. 3d,g).

Subsequently, we explored whether the gut microbiota composition in wild-type and *per*^01^ host genotypes could be modulated by the 12:12 LD and DD lighting conditions. When we measured microbiota alpha diversities, the species richness did not significantly vary between 12:12 LD and DD conditions in wild-type flies (Supplementary Fig. S4a; Chao1 index comparison: P > 0.05, ns). On the contrary, lower richness levels were found in *per*^01^ flies subjected to DD conditions compared to those maintained in 12:12 LD cycles (Supplementary Fig. S4b; Chao1 index comparison: P = 0.046). No variations in the Shannon index were observed for both genotypes in the two LD and DD lighting conditions (Supplementary Fig. S4a,b; P> 0.05, ns for both Canton-S and *per*^01^). Moreover, the beta diversity analysis revealed no differences in gut microbiota composition between 12:12 LD and DD regimes in both fly strains (Supplementary Fig. S4c,d; P> 0.05, ns for both Canton-S and *per*^01^).

### 2.3 Some components of the fly gut microbiota show daily variations which appear independent from the host’s circadian clock

We next asked whether wild-type and *per*^01^ microbiota displayed significant daily oscillations in 12:12 LD and/or DD conditions (Fig. 4a). We used alpha and beta diversity metrics to estimate possible daily changes in wild-type and *per^01^*microbiota structure and composition, in both 12:12 LD and DD regimes. In both host genotypes, total species richness and evenness remained stable throughout the day, in 12:12 LD as well as DD conditions (Fig. 4b-e; Chao1 and the Shannon indices: P > 0.05, ns for all comparisons). However, beta diversity analysis revealed that in wild-type guts the microbial composition significantly varied during the 24h day in 12:12 LD, while no significant changes were detected in DD conditions [Fig. 4f,g; in (f) 12:12 LD: P = 0.016; in (g) DD: P = 0.057, ns]. Surprisingly, microbiota of *per*^01^ mutants did not show any significant daily variations in composition in 12:12 LD regimes, but significant daily modifications in DD conditions [Fig. 4h,i; in (h) 12:12 LD: P = 0.09, ns; in (i) DD: P = 0.005) ].

**Figure 4:**
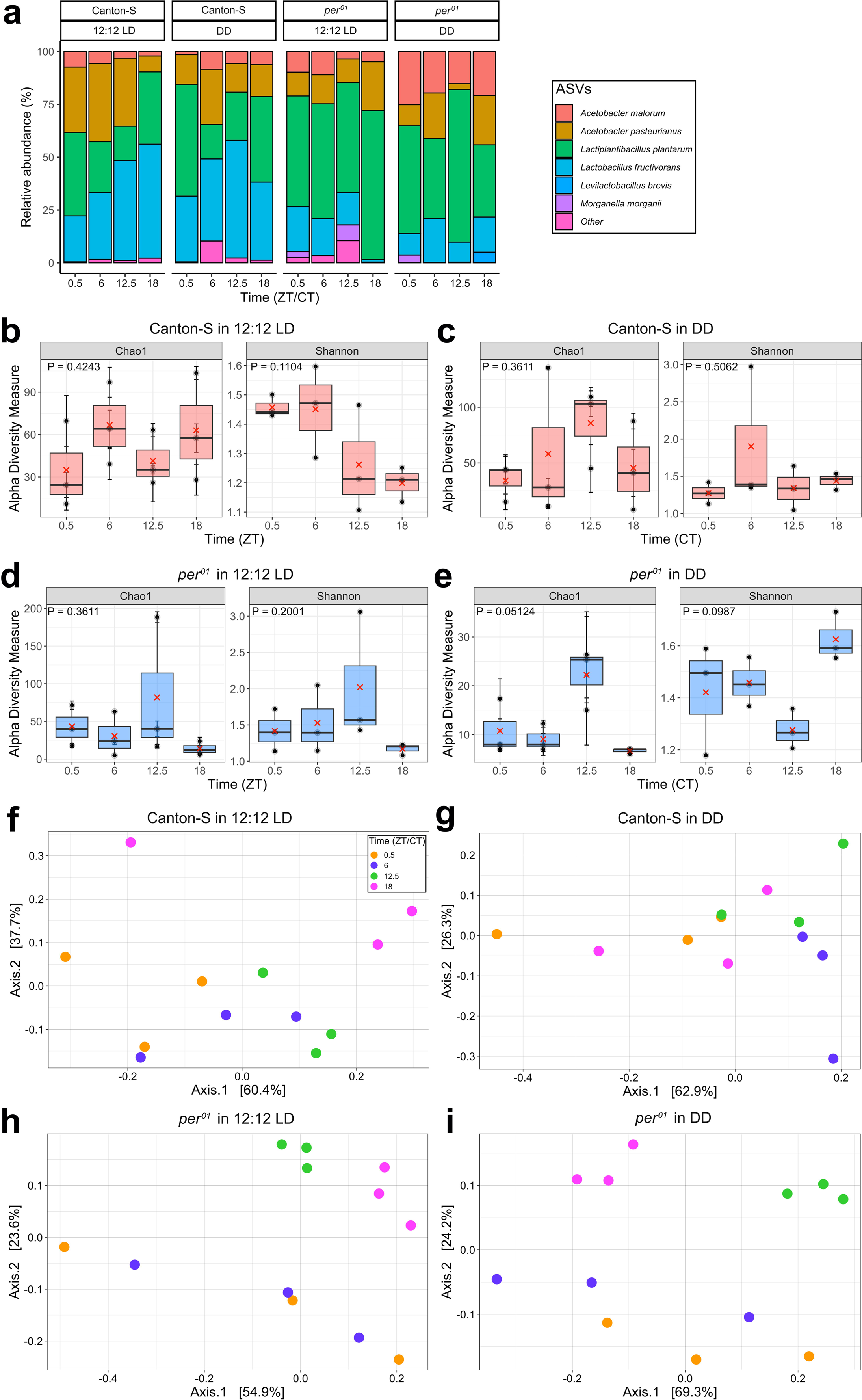
Gut microbiota daily variations in wild-type and *per^01^* flies under 12:12 LD and DD conditions. (a) Stacked bar chart showing the relative abundances (%) of the most prevalent bacterial ASVs in wild-type (Canton-S; eft) and *per^01^* (right) guts, at four different time points (ZTs/CTs 0.5, 6, 12.5, and 18) under 12:12 LD and DD conditions (**b, c**) Shannon and Chao1 alpha diversity indices of Canton-S microbiota, at four different time points (ZTs/CTs 0.5, 6, 12.5, and 18) under (**b**) 12:12 LD and (**c**) DD conditions. No significant daily differences in Chao1 and Shannon indices were detected in 12:12 LD and DD conditions [(**b**) 12:12 LD: Chao1: P = 0.424, ns; Shannon: P = 0.110, ns; (**c**) DD: Chao1: P = 0.361, ns; Shannon: P = 0.506, ns; Kruskal-Wallis test). (**d, e**) Shannon and Chao1 alpha diversity indices of *per^01^* microbiota at four different time points (ZTs/CTs 0.5, 6, 12.5, and 18) under (**d**) 12:12 LD and (**e**) DD conditions. No significant daily differences in Chao1 and Shannon indices were detected in 12:12 LD and DD conditions [(**d**) 12:12 LD: Chao1: P = 0.361, ns; Shannon: P = 0.200, ns; (**e**) DD: Chao1: P = 0.051, ns; Shannon: P = 0.099, ns; Kruskal-Wallis test). In each box plot, mean and median values are shown by red Xs and horizontal black lines, respectively. (**f, g**) PCoA of Bray-Curtis distances of Canton-S microbiota under (**f**) 12:12 LD and (**g**) DD regimes. PERMANOVA analyses showed significant differences in gut microbiota composition of Canton-S under (**f**) 12:12 LD (P = 0.016), but not in (**g**) DD (P = 0.057) conditions. (**h, i**) PCoA of Bray-Curtis distances of *per^01^*microbiota, in (**h**) 12:12 LD and (**i**) DD regimes. PERMANOVA analyses showed significant differences in gut microbiota composition of *per^01^* in (**i**) DD (P = 0.005), but not in (**h**) 12:12 LD (P = 0.09) conditions. Different symbols indicate different time points: circle = ZT/CT 0.5, cross = ZT/CT 6, triangle = ZT/CT 12.5, square = ZT/CT 18.

To explore these results, we first examined the 24 h abundance profiles of the dominant families *Lactobacillaceae* and *Acetobacteraceae* showing that they were not characterized by any significant cycling variation in both host genotypes in LD and DD regimes (Supplementary Fig. S5a-d; Supplementary Fig. S6a-d; P > 0.05, ns for all comparisons). Focusing on the most prevalent ASVs, we found that in wild-type guts, *A. malorum* was the unique ASV which exhibited a statistically significant variation, with a minimum during the night, in 12:12 LD regimes (Fig. 4a; Supplementary Fig. S5e-h; P< 0.05 only for *A. malorum* in Fig. S5h). Under DD conditions, in the same host genotype the *L. plantarum* ASV cluster displayed significant 24 h oscillations, with reduced levels at CT 6 and CT 12.5 (Fig. 4a; Supplementary Fig. S5i-l; P< 0.05 only for *L. plantarum* in Fig. S5i).

In the case of *per*^01^ flies in 12:12 LD regimes, no significant fluctuations in the relative levels of any ASVs were detected (Fig. 4a; Supplementary Fig. S6e-j; P> 0.05, ns for all comparisons), while only *A. pasteurianus* and *L. brevis* ASVs showed significant daily variations in DD conditions (Fig. 4a; Supplementary Fig. S6k-p; P< 0.05, for both *A. pasteurianus* and *L. brevis* in Fig. S6m and S6p, respectively). In particular, *A. pasteurianus* relative quantity increased every 12 h (at CTs 6 and 18), while *L. brevis* relative levels were low during the subjective day and increased at CT 18 in the subjective night (Fig. S6m,p).

### 2.4 Exploring the interplay between microbiota and feeding behavior in *Drosophila*

Feeding time is known to impact on the 24 h rhythmicity of the gut microbiota composition in mice and humans, ^13,34^. *D. melanogaster* was demonstrated to have a circadian rhythm in feeding behavior ^35^. Surprisingly this rhythmicity did not modulate the daily microbiota structure, in 12:12 LD conditions ^29^. However, different *Drosophila* strains can show different daily feeding profiles ^35^. To understand whether the variations in gut microbiome we detected in our study were in some way associated with feeding patterns in our *Drosophila* lines, we assessed the 24 h feeding behaviors of both wild-type and *per*^01^ flies kept under 12:12 LD and DD conditions using the Capillary Feeder Assay (CAFE) assay ^36^.

Wild-type flies exposed to a 12:12 LD cycle exhibited a rhythmic feeding behavior, with a first small but significant peak at ZT 2-4 and a second stronger one at ZT 10-12, (Fig. 5a, one-way ANOVA: P< 0.0001; JTK_CYCLE: P < 0.001). Under DD conditions, analysis of variance detected a significant daily variation in feeding behavior, with a small peak at CT 12-14 (Fig. 5b; one-way ANOVA: P<0.05), although the JTK_CYCLE algorithm did not show any significant circadian rhythmicity (P = 0.53, ns). Similar results were previously obtained using other wild-type strains and suggest that the fly feeding behavior has a weak/damped rhythmicity in DD, as reported by (Barber *et al.*, 2016).

**Figure 5:**
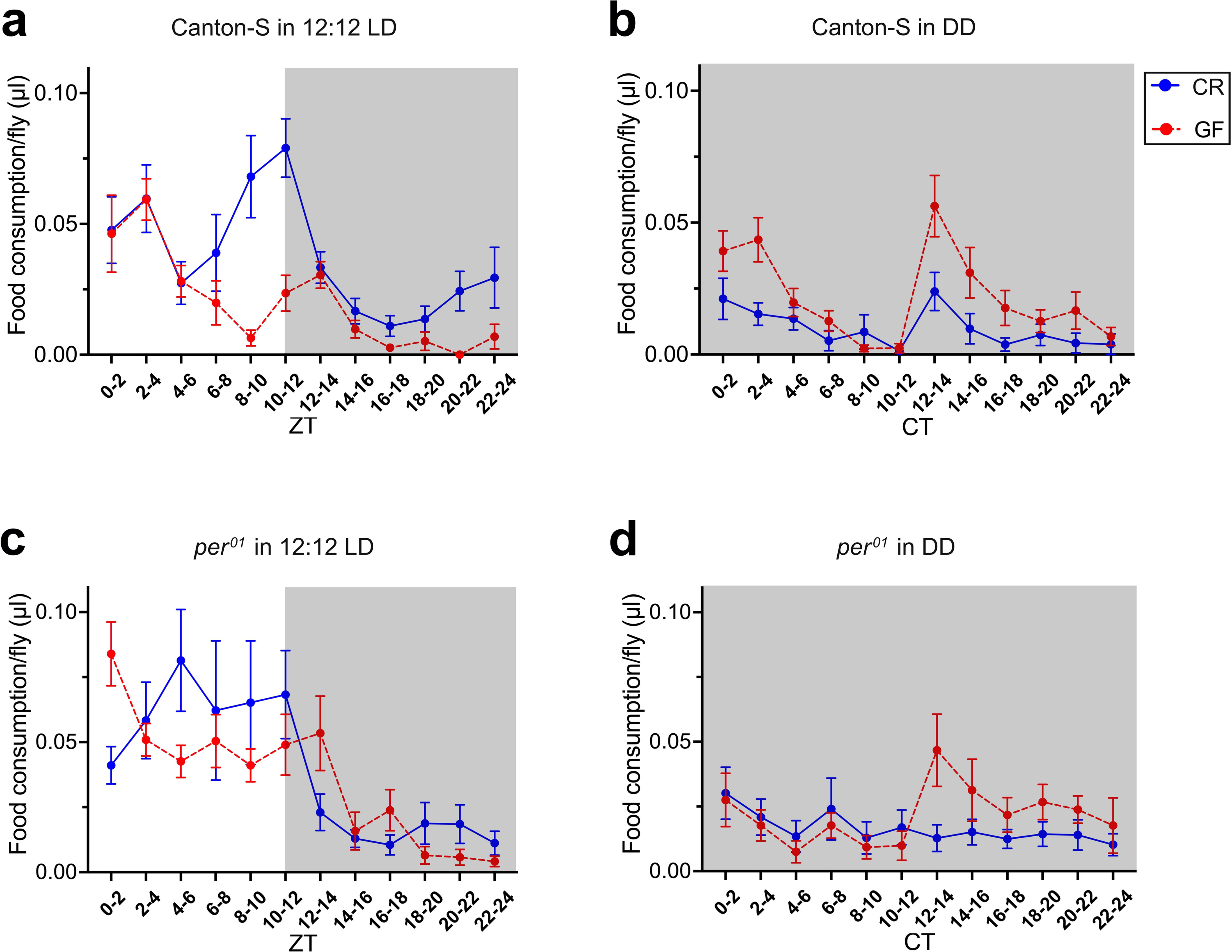
Daily feeding profiles of CR and GF wild-type and *per^01^* male flies under 12:12 LD and DD regimes. Each panel shows the amount of food intake per fly (mean μLs ± SEM) plotted against time, at 2-h intervals (ZTs/CTs). White and grey areas indicate the light and dark periods of the day, respectively. Feeding profiles are represented by a solid blue lines (CR flies), and dotted red lines (GF flies). (**a, b**) Daily feeding profiles of CR and GF Canton-S flies under (**a**) 12:12 LD and (**b**) DD regimes. Under (**a**) 12:12 LD cycles, both CR and GF wild-type (Canton-S) flies exhibited a robust rhythmic feeding pattern (CR: one-way ANOVA: P < 0.0001; JTK_CYCLE: P = 0.0002. Three independent experiments, N = 274. GF: one-way ANOVA: P < 0.0001; JTK_CYCLE: P < 0.0001. Two independent experiments, N = 141). Under (**b**) DD conditions, feeding behaviors of both CR and GF Canton-S flies showed significant daily variations with one-way ANOVA, but not with JTK_CYCLE algorithm (CR: one-way ANOVA: P = 0.024; JTK_CYCLE: P = 0.530, ns. Three independent experiments, N = 174; GF: one-way ANOVA: P < 0.0001; JTK_CYCLE: P = 1, ns. Two independent experiments, N = 114). (**c, d**) Daily feeding profiles of CR and GF *per^01^* flies under (**c**) 12:12 LD and (**d**) DD conditions. Under (**c**) 12:12 LD cycles, both CR and GF *per^01^* flies displayed strong rhythmicity (CR: one-way ANOVA: P = 0.0003; JTK_CYCLE: P < 0.0001. Four independent experiments, N = 262; GF: one-way ANOVA: P < 0.0001; JTK_CYCLE: P < 0.0001. Two independent experiments, N = 133). Under (**d**) DD conditions, both CR and GF *per^01^* exhibited an arrhythmic feeding profile (CR: one-way ANOVA: P = 0.7131, ns; JTK_CYCLE: P = 1, ns. Three independent experiments, N = 203; GF: one-way ANOVA: P = 0.0533, ns; JTK_CYCLE: P = 1, ns. Three independent experiments, N = 121).

*per*^01^ mutant flies exhibited a significantly increased food consumption during the light phase compared to the dark period of the day in 12.12 LD conditions (Fig. 5c; one-way ANOVA: P< 0.001; JTK_CYCLE: P < 0.0001). This outcome is likely due to a masking effect, where the presence of light induces higher food consumption even with a non-functional circadian clock, as in these clock mutant flies the rhythmicity in food intake was completely abolished in DD conditions (Fig. 5d; one-way ANOVA: P > 0.05, ns; JTK_CYCLE: P= 1, ns).

Finally, we asked whether gut bacteria exert an influence on host’s feeding patterns. To this end, we compared the feeding profiles of wild-type and *per*^01^ flies kept in germ-free (GF) conditions to those obtained from their counterparts colonized by gut microbes, named conventionally reared (CR) flies, under both 12:12 LD and DD conditions.

GF wild-type males showed a significant daily variation in feeding activity in 12:12 LD, although with slight modifications in the profile compared to CR wild-type flies (Fig. 5a; one-way ANOVA: P < 0.0001; JTK_CYCLE: P **<** 0.0001). Specifically, although the peak observed in CR flies at ZT 2-4 was maintained, the second peak at ZT10-12 detected in CR individuals occurred two hours later and was smaller in axenic conditions. GF wild-type flies showed feeding patterns similar to CR individuals in DD conditions, displaying a weak rhythmicity (Fig. 5b; one-way ANOVA: P < 0.0001; JTK_CYCLE: P = 1, ns).

In GF *per*^01^ flies, the food consumption was higher during the light phase compared to the dark phase under 12:12 LD conditions (Fig. 5c; one-way ANOVA: P< 0.0001; JTK_CYCLE: P < 0.0001), as previously observed for CR *per*^01^ mutants. In DD conditions, rhythmicity was also lost in GF *per*^01^ flies (Fig. 5d; one-way ANOVA: P> 0.05, ns; JTK_CYCLE *P* = 1, ns), and their feeding profile was similar to CR flies.

## 3. Discussion

In this study, we analyzed the gut microbiota composition and abundance in wild-type flies compared to isogenic *per^01^* arrhythmic mutants. Flies were reared under both 12:12 LD and DD conditions while being fed *ad libitum*. Our objective was to uncover potential links between the gut microbiota and the circadian clock in *Drosophila*, to advance our understanding of this relationship.

Wild-type and *per^01^* flies possessed a conventional microbiota characterized by most of the typical *Drosophila* gut bacteria belonging to *Acetobacteraceae* and *Lactobacillaceae* families ^26,37^. However, wild-type and *per^01^* flies showed different microbiota compositions in both 12:12 LD and DD conditions, as revealed by beta diversity analyses. These dissimilarities might be explained by the presence of ASVs specific to one of the two host genotypes, as well as by variations in relative levels of ASVs common to both wild-type and *per^01^* microbiota. In fact, when we analyzed the most prevalent ASVs, *M. morganii* and *L. brevis* were exclusively found in *per^01^*flies. Among the shared ASVs, *per^01^* microbiota showed significantly lower relative abundances of *L. fructivorans* and *A. pasteurianus* and higher relative quantities of *A. malorum* and *L. plantarum* compared to wild-type flies. The wild-type and *per^01^*strains used in the present study have been consistently reared in our laboratory under comparable conditions for several years. Both lines were fed the same diet and underwent parallel processing for the microbiota analyses. Thus, it is unlikely that the observed differences in microbiota between the two host genotypes are due to stochastic factors, which have been reported to contribute in determining microbiota composition in *Drosophila* ^38^. Taken together, these results indicate that the circadian clock significantly impacts the composition of the fly gut microbiota. Interestingly, a functional clock seems also important in maintaining a stable microbial species richness in the absence of any LD cycle cue, as the gut microbiota of *per^01^*flies maintained in DD showed a significant lower total species richness (Chao1 index) compared to those of both *per^01^* in 12:12 LD and wild-type flies in 12:12 LD or DD regimes. Extending these analyses to other fly strains carrying mutations at the level of other cardinal clock genes in a cantonized genetic background, and using gnotobiotic lines will clarify these aspects. However, it is interesting to mention that gut microbial composition was significantly affected in mouse when the circadian clock was genetically disrupted in the whole organism (*Per1/2* ^-/-^ double mutant) or specifically in intestinal enterocytic cells (epithelial cell-driven knock-out of the core clock gene *Bmal1*) ^13,17^. Collectively, these data suggest that the circadian clock influences microbiome composition in both *Drosophila* and mammals.

When we searched for daily variations in gut microbiota total abundances, we showed that the overall amount of bacteria was stable across the day in wild-type flies under both 12:12 LD and DD conditions. In contrast, *per^01^* mutants exhibited significant 24 h variations in gut bacterial levels, with total bacterial abundance decreasing by 15–20 % during nighttime (ZT/CT18) under 12:12 LD and DD regimes. Interestingly, gut microbiota abundance did not correlate with the daily feeding behavior, which has been demonstrated to be under circadian control ^39^. In fact, wild type flies, which did not show any change in bacterial levels under both 12:12 LD and DD conditions, displayed a clear rhythmic bimodal feeding profile in 12: 12 LD and a unimodal profile with a damped rhythmicity under DD conditions. In contrast, *per^01^* mutant flies showed decreasing microbiota loads during nighttime, regardless of whether they exhibited rhythmic (12:12 LD) or erratic and arrhythmic (DD) feeding behaviors. Additionally, conventionally reared wild-type and *per^01^*flies maintained substantially comparable feeding profiles in germ free conditions, under both 12: 12 LD and DD regimes. These results mirrored those previously obtained using other isogenic wild-type and *per^01^*lines ^29^, characterized by a different genetic background with respect to our strains, and indicate that the gut microbiota does not affect the daily feeding profile in *Drosophila melanogaster*.

When we looked for daily modifications in the gut microbiota with-in structures of wild-type and *per*^01^ flies using alpha diversity analyses, we found that neither wild-type nor *per^01^* microbiota showed significant variations under 12:12 LD and DD conditions. These results paralleled the absence of daily fluctuations in species richness and evenness previously reported in microbiota derived from fecal samples of wild-type and *per^01^* flies maintained under 12:12 LD regimes and fed *ad libitum* (Zhang et al 2023). The same authors detected a limited daily fluctuation in microbiota species richness and evenness only in *per^01^* flies, when fed under a TF paradigm, a condition known to promote cycling ^29^.

Taken together, these results suggest that a functional circadian clock plays a role mainly in stabilizing gut microbial abundances over the course of the day. Several data indicate that the circadian clock has an important function in guaranteeing daily fluctuations in the composition of the intestinal microbiota in both mice and humans ^13,14,17^. On the contrary, it seems that the fly circadian clock ensures stability of the microbiota abundance (this work) and structure only in particular conditions, such as TF (as reported in ^29^).

Finally, our beta diversity analyses suggest the existence of significant variations in gut microbiota daily composition of wild-type and *per^01^*flies, which appeared independent from a functional circadian clock. In fact, significant dissimilarities in microbiota daily composition were detected in wild-type flies but not in *per^01^* mutants under 12:12 LD conditions, and in *per^01^* mutants but not in wild-type individuals under DD regimes. Although further analyses will clarify these results, it is interesting to mention that some of the most prevalent ASVs showed significant daily fluctuations in their relative abundance in both wild type and *per^01^* genotypes (i.e., in wild type: *A. malorum* under 12:12 LD, and *L. plantarum* under DD conditions; in *per^01^: A. pasteurianus* and *L. brevis*, under DD regimes). These ASVs exhibited different profiles of variation, with certain ASVs showing higher relative levels during the day (e.g., *A. malorum* in wild-type guts under 12:12 LD), while others during the night (e.g., *L. plantarum* in wild-type guts under DD). Moreover, these variations did not appear to be correlated with the feeding profile. For instance, significant daily variations in *A. pasteurianus* and *L. brevis* ASVs were detected in *per^01^* flies under DD, a condition determining an arrhythmic feeding behavior in this clock mutant. These results might indicate that certain gut commensal bacteria in D*. melanogaster* have an intrinsic circadian clock that operates independently of the host’s circadian rhythms. Procaryotic circadian clocks have been well documented in cyanobacteria ^40–43^ and recently reported in the free-living non-photosynthetic bacterium *Bacillus subtilis* when growing as biofilms ^44^. Importantly, it was shown that the enteric bacterium *Enterobacter aerogens* (i.e., *Klebsiella aerogens*) possesses a circadian rhythm in swarming motility, which can be synchronized by temperature and melatonin stimuli, indicating that at least one member of the human microbiome has an endogenous circadian clock which might be entrained by host’s circadian cues ^45,46^. Additional studies are warranted to determine the presence and functionality of circadian mechanisms within the gut commensal bacteria of *D. melanogaster*. Furthermore, exploring whether these mechanisms contribute to shaping the daily intra- and inter-species dynamics of the gut microbiome would provide valuable insights into the link between the host circadian system and its associated microbial community.

In summary, our data suggest that the *Drosophila melanogaster* circadian clock has a role in shaping the gut microbiome composition, as demonstrated in mammals. Additionally, a functional clock appears to prevent daily oscillations in the gut bacterial total abundance, suggesting that in *D. melanogaster* the circadian clock is important for guarantying a daily stability of the gut microbiome load. Surprisingly, our analyses also indicated that a few components of the fly gut microbiota present significant daily fluctuations in their relative abundance, which appeared independent from the host’s circadian clock. Future investigations are required to understand the potential benefit of a constant daily microbiome load for fly fitness and to elucidate whether in *Drosophila* some gut commensal bacteria have a host-independent circadian clock.

## 4. Materials and Methods

### Fly strains and maintenance

The *D. melanogaster* strains used in this study were: Canton-S and the cantonized clock mutant line *per^01^* (University of Leicester, Leicester, UK; ^30^). Fly stocks were routinely maintained in 12:12 LD regime (850 lux) at 23°C, on a cornmeal standard diet (7.2 % cornmeal, 7.9 % sucrose, 5 % dried yeast, 0.85 % agar, 0.3% propionic acid, and 0.27 % nipagin).

### PCR control of the *per^+^* and *per^01^* alleles and detection of possible *Wolbachia spp*. contaminations in Canton-S and *per^01^* strains.

See Supplementary information.

#### Analysis of locomotor activity behavior

See Supplementary information.

### Gut sample collection and DNA extraction for microbiota analysis

Environmental factors can affect gut microbiome compositions. To avoid gut microbiome variations due to undesired environmental variables, Canton-S and *per^01^* strains were reared in parallel on the same batches of food at 23 °C in 12:12 LD conditions, for at least one month before sampling. During this period, flies were transferred into fresh food-containing vials every 2-3 days. At the beginning of the experiment, flies were either kept under 12:12 LD conditions or entrained for three days under 12:12 LD and then transferred for two days under DD. Four-to-seven-day-old adults were collected at four different time points (ZTs/CTs 0.5, 6, 12.5, 18) in 12:12 LD or on the third day of DD. Each sampling was performed transferring anesthetized flies into a collection basket (Biosigma), equipped with a 100 μm nylon mesh (Biosigma) and covered with sterile gauze. To remove external bacteria, the basket was submerged 30 s in 70% ethanol and washed thrice in sterile PBS. Flies were transferred into new tubes, left 3 min in a dry ice-ethanol bath, and then stored at -80°C until processing. Guts were dissected only from males in PBS (pH 7.4), using sterilized tools. Gut samples included cardia, crop, foregut, midgut, and hindgut, whereas Malpighian tubules were excluded. For each time point, three replicates with 6-20 guts were collected in a screw-cap tube containing 300 μl of DNA/RNA Shield (Zymo Research) (details on number of guts obtained for each sample can be found in Supplementary Table S1).

For microbial community analysis, three negative controls (mocks) were prepared by pipetting in a tube 100 μl of PBS in which analogous handlings for dissections (without flies) were performed.

Genomic DNA (gDNA) extraction was performed under sterile conditions using the Qiagen Blood & Tissue Kit (Qiagen) with a lysozyme pre-treatment, following the manufacturer’s instructions. Briefly, dissected guts were washed with sterile PBS, homogenized 1 min in a TeSeE PRECESS 24 (Bio-Rad) using 0.5 mm glass beads (Sigma-Aldrich) in 180 μl of an enzymatic lysis buffer containing 20 mM Tris HCl at pH 8.0, 2 mM sodium EDTA, 1.2 % Triton X-100, and 20 mg/mL lysozyme (Sigma-Aldrich). Homogenates were incubated 1.5 h at 37 °C. Each supernatant was transferred into a new tube and incubated with 25 μL Proteinase K 30 min at 56 °C. After addition of 200 μL absolute ethanol, each sample was transferred to DNeasy Mini spin column. Columns were centrifuged, washed, and DNA was eluted in 50 μL RNase/DNase-free water.

To assess the efficiency in bacterial DNA extraction from gut samples, DNA from the ZymoBIOMICS™ Microbial Community Standard (Zymo Research) was processed in parallel as a positive control (Supplementary Table S2). To identify possible contaminants deriving from gut dissection procedures, DNA was extracted from the three mocks (negative controls), collected during dissections.

### Estimation of total microbiota abundance

The total abundance of gut microbiota was assessed via qPCR using the following universal primers F: 5’-TCCTACGGGAGGCAGCAGT-3’ and R: 5’-GGACTACCAGGGTATCTAATCCTGTT-3’, targeting the V3-V4 regions of the *16S* rDNA gene ^47^. *Drosophila rp49* was used to normalize the amount of detected *16S* rDNAs, using the primers 341F: 5’-GCCGCTTCAAGGGACAGTATCTG-3’ and 805R: 5’-AAACGCGGTTCTGCATGA-3’.

For each sample, reactions were prepared in triplicate, using TB Green *Premix Ex Taq* (TaKaRa) and 0.2 μM of each primer. Amplification cycling conditions were: pre-incubation at 95 °C 30 s, 40 cycles at 95 °C 5 s and 60 °C 30 s. qPCR was performed on a CFX96 Real-Time PCR System (BioRad). Analysis of melting curves confirmed amplification specificity. Primer efficiencies were determined using a series of four 10-fold dilutions from random samples, resulting in 91.15 % and 92.11 % for 16S rDNA and *rp49*, respectively. Data were expressed as a ratio between Ct values of *16S* and *rp49* genes.

### Sequencing, read processing and microbial diversity analyses

Illumina sequencing targeting V3-V4 regions of the *16S* rDNA gene was performed by the IGATech sequencing center (IGA Technology Services s.r.l., Udine, Italy) on an Illumina MiSeq platform (Illumina, San Diego, CA) using 300-bp paired-end mode. A total of 2,951,031 reads were obtained from all samples, with a mean of 56,750.596 reads per sample. The range of reads per sample varied from a minimum of 14,255 to a maximum of 92,186. The number of reads obtained for each sample can be found in Supplementary Table S1.

Raw data quality was inspected using FastQC (version 0.11.9) ^48^. Reads were analyzed using QIIME 2 (version 2022.11) ^49^. In particular, the DADA2 plugin (version 1.26.0) ^50^ was used to merge forward and reverse reads, identify chimeras (“consensus” method), and trim low quality portion of reads (trunc-len-f = 280, trunc-len-r = 220). ASV taxonomy was assigned using the feature-classifier and classify-sklearn method on the SILVA reference database trained on the *16S* rDNA portion amplified by the 341F-805R primers ^51^. ASVs sequences were processed using phylogeny align-to-tree-mafft-fasttree plugin for multiple alignments (MAFFT software, version 7.508) ^52^, and then unrooted and rooted maximum-likelihood phylogenetic trees were inferred with FastTree (version 2.1.11) ^53^. The total number of ASVs identified in all samples was 4351. Principal components analysis (PCoA) performed on raw data confirmed that mocks significantly differed from the other samples (Supplementary Figure S7). This indicates that the experimental protocol successfully extracted and sequenced gut bacterial DNA rather than environmental contaminants. Before any additional analysis, ASVs were subjected to three filtering steps. Potential contaminants deriving from dissection steps were identified using the *decontam* package (version 1.20.0) ^54^ in R (version 2023.03.1+446) based on the frequencies of ASVs in the three negative controls using default parameters with a probability threshold set at 0.5. 43 ASVs were identified as contaminants and removed at this step. Secondly, ASVs that were annotated as chloroplasts or mitochondria were removed (15 ASVs). Lastly, ASV sequences shorter than 400 bp were removed. These sequences had no hit in the SILVA database and were classified as “unknown”. After a blastn search in the nr NCBI database, they were identified as portions of the *16S* gene of chloroplasts or mitochondria from plants or *Drosophila*. Therefore, these short ASVs were dropped from further processing. At this final step, 2417 ASVs were removed from the original dataset. The total number of ASVs removed after these three filtering steps was 2432, leaving 1919 ASVs that were retained for the following analysis. Data were normalized using the rarefy_even_depth function in the R package Phyloseq (version 1.44.0) ^55^ to 90% of the smallest sample size (PLB2 sample code), and subsequent analyses on microbial composition were performed on the rarefied data set. Quantification of alpha diversity was performed using the estimate_richness function in Phyloseq using Shannon diversity and Chao1 indices. Beta diversity was estimated based on Bray-Curtis dissimilarity with the distance function in Phyloseq, and visualized via Principal Coordinate Analysis (PCoA) ordination.

### Generation and rearing of GF flies

GF flies were generated as in ^56^ with slight modifications. Briefly, flies were transferred in a cage (Biosigma) covered with a fruit juice agar plate (24.8 % apple or peach juice, 2.48 % sucrose, 3 % agar), and a small amount of yeast paste placed at the center (ratio 9:1 brewer’s dry yeast: H_2_O). Flies were left to lay eggs for about 16 h before embryo collection. Afterwards, the surface of the agar plate was covered with sterile water and gently brushed to detach embryos from the agar. Embryos were collected inside a laminar flow hood, using a basket (Biosigma) equipped with a 100 μm nylon mesh (Biosigma), sterilized for 2 min with 70 % ethanol followed by a 10 min treatment with 10 % bleach, and washed three times in sterile ddH_2_O. Dechorionated eggs were then transferred into vials filled with a sterile diet. The food for GF flies was prepared by autoclaving an 1.8 % agar-supplemented cornmeal standard diet. After cooling at 50 °C, nipagin and propionic acid were added to the autoclaved diet. The sterile condition of GF flies was periodically tested plating a homogenate of flies on a Lysogeny Broth (LB) agar plate.

### CAFE assay

The feeding assay was performed as in ^36^, with slight modifications. To familiarize with the food source, one day prior to the experiment, six 2-9 day-old males were placed into a standard plastic vial (Greiner Bio-one) containing a 4 X 4 cm filter paper soaked with 500 μl ddH_2_O, and four 5 μl calibrated capillaries (BLAUBRAND® micropipettes), filled with a 5% sucrose solution in ddH_2_O. For each replicate, six fly-containing vials were placed in a plastic box together with two plastic vials, containing 50 mL ddH_2_O and left open to ensure proper humidity. Additionally, three vials without flies were placed in the same box to estimate evaporation rates during the experiment. On the day of the experiment, the capillaries containing the sucrose solution were discarded from each vial and replaced with capillaries filled with a 5% sucrose and 0.25 % patent blue E131 (Fabbrica Italiana Coloranti per Alimenti) solution in ddH_2_O. The meniscus levels of the capillaries were marked every two hours, then length measurements (in mm) were converted into volumes (µL) and divided for the number of flies in each vial, considering the average amount of evaporation. All feeding assays were performed in a temperature-controlled incubator at 23 °C. For experiments under DD conditions, flies were entrained for at least 3 days to 12:12 LD condition before transferring to DD. The feeding curves were registered on the third day in DD. Experiments on GF flies were performed under sterile conditions, using autoclaved or UV-treated materials.

### Statistical analyses

QPCR data on total bacterial abundance, alpha diversity Chao and Shannon indices, and single taxa daily changes in relative abundances did not approximate normal distributions, evaluated with Shapiro-Wilk test. Thus, they were non-parametrically analyzed using Kruskal-Wallis test, followed by Dunn’s *post hoc* test.

A permutational multivariate analysis of variance (PERMANOVA) implemented in the adonis2 function of the Vegan package (version 2.6.4) was used to analyze beta diversity measures ^57^. Two-tailed t-tests were used to compare the relative abundance of the prevalent bacterial families and ASVs between Canton-S and *per^01^*microbiota. Details on numbers of dissected guts, replicates, and sequence reads obtained for microbiota data generation are reported in Supplementary Table S1. Sequencing data are available at NCBI BioProject with accession PRJNA1087739. Feeding profiles were analyzed using the ordinary one-way ANOVA followed by Tukey’s multiple comparison test, and the rhythmicity was tested with the JTK_CYCLE algorithm (version 3.1) ^58^. Two to four independent experiments were performed for feeding profiles determination, with N ranging from 114 to 274 individuals. Statistical analyses were conducted in R (version 2023.12.0) or in GraphPad Prism (version 9.5.1).

#### Author contribution

Conceived and designed the experiments: OR, FS, MB. Performed the experiments: MB, IV. Analyzed the data: MB, OR. Contributed reagents/materials/ analysis tools: FS. Wrote the manuscript: MB, OR, FS.

#### Competing interests

The authors have declared that no competing interests exist

#### Funding

This work was funded by Cinchron, a European Union’s Horizon 2020 research and innovation programme under the Marie Sklodowska-Curie (grant agreement No 765937) to FS. IV was supported by a fellowship from the Department of Biology, Università degli Studi di Padova (LANF_ECCELLENZA18_01). OR was supported by a Roux-Pasteur-Cantarini postdoctoral fellowship. The funders had no role in study design, data collection and analysis, decision to publish, or preparation of the manuscript.

## Acknowledgements

We are grateful to the *Drosophila* facility at the Department of Biology, Università degli Studi di Padova for *Drosophila* lines maintenance.

## Notes

### Competing Interest Statement

The authors have declared no competing interest.

