## Supplementary for "A functional circadian clock regulates composition and daily bacterial load of the gut microbiome in *Drosophila melanogaster*"

### Supplementary information

Matteo Battistolli[1], Irene Varponi[1], Ottavia Romoli[2]*, Federica Sandrelli[1]*

[1] Department of Biology, University of Padova, Padova, Italy

[2] Institut Pasteur, Université Paris Cité, CNRS UMR3569, Viruses and RNAi, F-75015 Paris, France

* Co-corresponding authors

### Supplementary Material and Methods

#### PCR control of the *per*$^{+}$ and *per*$^{01}$ alleles and detection of possible *Wolbachia* spp. contaminations in Canton-S and *per*$^{01}$ strains

Genomic DNA (gDNA) was obtained from a pool of five homogenized flies, using Wizard® Genomic DNA purification Kit (Promega) following the manufacturer's instructions. Briefly, flies were ground using a pestle in 0.5 mL dd$H_2O$ and centrifuged 2 min at 14,000 rpm. 600 µL Nuclei Lysis Solution were added to each pellet and samples were incubated 5 min at 80 °C. Samples were then incubated with 3 µL RNase 45 min at 37 °C and cooled at room temperature. After addition of 200 µL Protein Precipitation Solution, samples were incubated 5 min on ice and centrifuged 3 min at 14,000 rpm. DNA samples were then precipitated with isopropanol, washed with 70% ethanol, re-suspended in DNA Rehydration Solution, and incubated 1 h at 65 °C. DNAs were stored at -20°C until use. The molecular discrimination between *per*$^{+}$ and *per*$^{01}$ alleles is possible because of the C->T substitution at nucleotide position 4,179 in the *per*$^{01}$ allele (NC_004354.4), which generates in the surrounding region a *Xba*I restriction site in *per*$^{01}$sequence[1]. *per* alleles were checked in wild-type (Canton-S) and *per*$^{01}$ strains via PCR, using the following primers: *perDm* Forward: 5'-CGCAG-CATCATGGACTTCTA-3' and *perDm* Reverse: 5'-GACGGTCAACTCGTTCTCGT-3', to obtain a 551 bp DNA fragment. Amplifications were performed in a final volume of 25 µL, containing 2X Dream Taq Green PCR Master Mix (ThermoFisher), 0.4 µM of each primer, and 2 ng of gDNA. The amplification cycle was: 95 °C 3 min, (95 °C 30 s, 64 °C 45 s, 72 °C 45 s) for 35 cycles, 10 min at 72 °C. Amplicons were digested with *Xba*I (New England Biolabs) 20 min at 37 °C, obtaining two different fragments (125 and 426 bp) in the case of *per*$^{01}$ and an uncut 551 bp segment, in the case of *per*$^{+}$ allele.

Detection of bacterial endosymbiont *Wolbachia* spp. was performed via amplification of *Wolbachia* surface protein (*wsp*) gene using the following primers *wsp* 81F: 5'-TGGTCCAA-TAAGTGATGAAGAAAC-3' and *wsp* 691R: 5'-AAAAATTAAACGCTACTCCA-3', as in [2].

Amplified fragments can vary from 590 to 632 bp, depending on the *Wolbachia* strain. Amplifications were performed in a final volume of 25 µL, with 2X Dream Taq Green PCR Master Mix (ThermoFisher), 0.4 µM of each primer, and 4 ng of gDNA, and with the following PCR cycle: 94 °C 3 min, (94 °C 1 min, 55 °C 1 min, 72 °C 1 min) for 35 cycles, and 72 °C 10 min. gDNA of known *Wolbachia*-infected flies was used as a positive control.

**Analysis of locomotor activity behavior**

Locomotor activity of 2–5-day-old males was recorded as 1 min bins for three days in 12:12 LD, followed by seven days under DD conditions, using the TriKinetics *Drosophila* Activity Monitor (DAMSystem2). Data were analyzed using a freely available Faas (Fly activity analysis suite) software (version 1.2) by Michel Boudinot and François Rouyer (https://neuropsi.cnrs.fr/en/departments/cnn/group-leader-francois-rouyer/).

**Supplementary Table S1: Details on sequenced microbiota samples. For each sample, host genotype, lighting condition, sampling time, progressive replicate number, number of dissected guts, and number of reads are indicated.**

| Sample * | Genotype | Lighting condition | Sampling time | Replicate | Number of guts per replicate ** | Reads number |
|---|---|---|---|---|---|---|
| CLA1 | Canton-S | 12:12 LD | ZT 0.5 | 1 | 20 | 59,570 |
| CLA2 | Canton-S | 12:12 LD | ZT 0.5 | 2 | 20 | 50,612 |
| CLA3 | Canton-S | 12:12 LD | ZT 0.5 | 3 | 20 | 62,662 |
| CLB1 | Canton-S | 12:12 LD | ZT 6 | 1 | 20 | 83,211 |
| CLB2 | Canton-S | 12:12 LD | ZT 6 | 2 | 20 | 52,616 |
| CLB3 | Canton-S | 12:12 LD | ZT 6 | 3 | 20 | 50,423 |
| CLC1 | Canton-S | 12:12 LD | ZT 12.5 | 1 | 7 | 51,497 |
| CLC2 | Canton-S | 12:12 LD | ZT 12.5 | 2 | 7 | 58,424 |
| CLC3 | Canton-S | 12:12 LD | ZT 12.5 | 3 | 6 | 50,871 |
| CLD1 | Canton-S | 12:12 LD | ZT 18 | 1 | 18 | 50,432 |
| CLD2 | Canton-S | 12:12 LD | ZT 18 | 2 | 17 | 92,186 |
| CLD3 | Canton-S | 12:12 LD | ZT 18 | 3 | 16 | 70,797 |
| CDA1 | Canton-S | DD | CT 0.5 | 1 | 20 | 58,769 |
| CDA2 | Canton-S | DD | CT 0.5 | 2 | 20 | 51,398 |
| CDA3 | Canton-S | DD | CT 0.5 | 3 | 20 | 62,440 |
| CDB1 | Canton-S | DD | CT 6 | 1 | 17 | 53,947 |
| CDB2 | Canton-S | DD | CT 6 | 2 | 17 | 49,525 |
| CDB3 | Canton-S | DD | CT 6 | 3 | 17 | 53,774 |
| CDC1 | Canton-S | DD | CT 12.5 | 1 | 16 | 57,217 |
| CDC2 | Canton-S | DD | CT 12.5 | 2 | 16 | 50,142 |
| CDC3 | Canton-S | DD | CT 12.5 | 3 | 16 | 52,048 |
| CDD1 | Canton-S | DD | CT 18 | 1 | 20 | 52,219 |
| CDD2 | Canton-S | DD | CT 18 | 2 | 20 | 56,383 |
| CDD3 | Canton-S | DD | CT 18 | 3 | 20 | 53,130 |
| PLA1 | $per^{01}$ | 12:12 LD | ZT 0.5 | 1 | 20 | 59,546 |
| PLA2 | $per^{01}$ | 12:12 LD | ZT 0.5 | 2 | 20 | 50,270 |
| PLA3 | $per^{01}$ | 12:12 LD | ZT 0.5 | 3 | 20 | 56,467 |
| PLB1 | $per^{01}$ | 12:12 LD | ZT 6 | 1 | 20 | 54,272 |
| PLB2 | $per^{01}$ | 12:12 LD | ZT 6 | 2 | 20 | 60,859 |
| PLB3 | $per^{01}$ | 12:12 LD | ZT 6 | 3 | 20 | 41,031 |
| PLC1 | $per^{01}$ | 12:12 LD | ZT 12.5 | 1 | 20 | 55,346 |
| PLC2 | $per^{01}$ | 12:12 LD | ZT 12.5 | 2 | 20 | 53,455 |
| PLC3 | $per^{01}$ | 12:12 LD | ZT 12.5 | 3 | 20 | 44,240 |
| PLD1 | $per^{01}$ | 12:12 LD | ZT 18 | 1 | 20 | 53,138 |
| PLD2 | $per^{01}$ | 12:12 LD | ZT 18 | 2 | 20 | 72,759 |
| PLD3 | $per^{01}$ | 12:12 LD | ZT 18 | 3 | 20 | 87,827 |
| PDA1 | $per^{01}$ | DD | CT 0.5 | 1 | 20 | 50,150 |
| PDA2 | $per^{01}$ | DD | CT 0.5 | 2 | 20 | 50,650 |
| PDA3 | $per^{01}$ | DD | CT 0.5 | 3 | 20 | 52,143 |
| PDB1 | $per^{01}$ | DD | CT 6 | 1 | 20 | 73,969 |
| PDB2 | $per^{01}$ | DD | CT 6 | 2 | 20 | 62,364 |
| PDB3 | $per^{01}$ | DD | CT 6 | 3 | 20 | 53,155 |
| PDC1 | $per^{01}$ | DD | CT 12.5 | 1 | 20 | 53,536 |
| PDC2 | $per^{01}$ | DD | CT 12.5 | 2 | 20 | 50,642 |
| PDC3 | $per^{01}$ | DD | CT 12.5 | 3 | 20 | 50,499 |
| PDD1 | $per^{01}$ | DD | CT 18 | 1 | 20 | 50,842 |
| PDD2 | $per^{01}$ | DD | CT 18 | 2 | 20 | 51,360 |
| PDD3 | $per^{01}$ | DD | CT 18 | 3 | 20 | 58,929 |
| MCK1 | N/A | N/A | N/A | N/A | N/A | 56,668 |
| MCK2 | N/A | N/A | N/A | N/A | N/A | 14,255 |
| MCK3 | N/A | N/A | N/A | N/A | N/A | 56,386 |
| CNTR+ | N/A | N/A | N/A | N/A | N/A | 91,980 |

* Sample code: first letter: "C" , Canton-S and "P", $per^{01}$ genotypes; second letter:"L", 12:12 LD and "D", DD conditions; third letter: "A": ZT/CT 0.5, "B":ZT/CT 6, "C":ZT/CT 12.5, and "D":ZT/CT 18. "MCK": negative control; "CNTR+" positive control. **: All samples derived from 16-20 dissected guts, with the exception of CLC1-3 replicates, each derived from 6-7 guts. However, these samples reached the plateau in rarefaction curves for both observed ASVs and Shannon index, indicating that their total microbial diversity was sequenced.

**Supplementary Table S2: Theoretical and obtained relative abundances (%) of Zymo-BIOMICS microbial community standard (Zymo Research).**

| Species | Gram-type | Theoretical relative abundance (%) | Obtained relative abundance (%) * |
|---|---|---|---|
| *Pseudomonas aeruginosa* | G- | 4.3 | 3.6 |
| *Escherichia coli* | G- | 10.1 | 19 |
| *Salmonella enterica* | G- | 10.4 | 15.8 |
| *Lactobacillus fermentum* | G+ | 18.4 | 8 |
| *Enterococcus faecalis* | G+ | 9.9 | 9.11 |
| *Staphylococcus aureus* | G+ | 15.5 | 12.8 |
| *Listeria monocytogenes* | G+ | 14.1 | 6.5 |
| *Bacillus subtilis* | G+ | 17.4 | 24 |
| *Saccharomyces cerevisiae* | NA | NA | NA |
| *Cryptococcus neoformans* | NA | NA | NA |

*Deviation of obtained from theoretical values ranged from 0.7 to-10.4 %. All microbial taxa were detected, indicating that DNA extraction and sequencing method used were adequate to detect multiple Gram positive and negative bacteria.

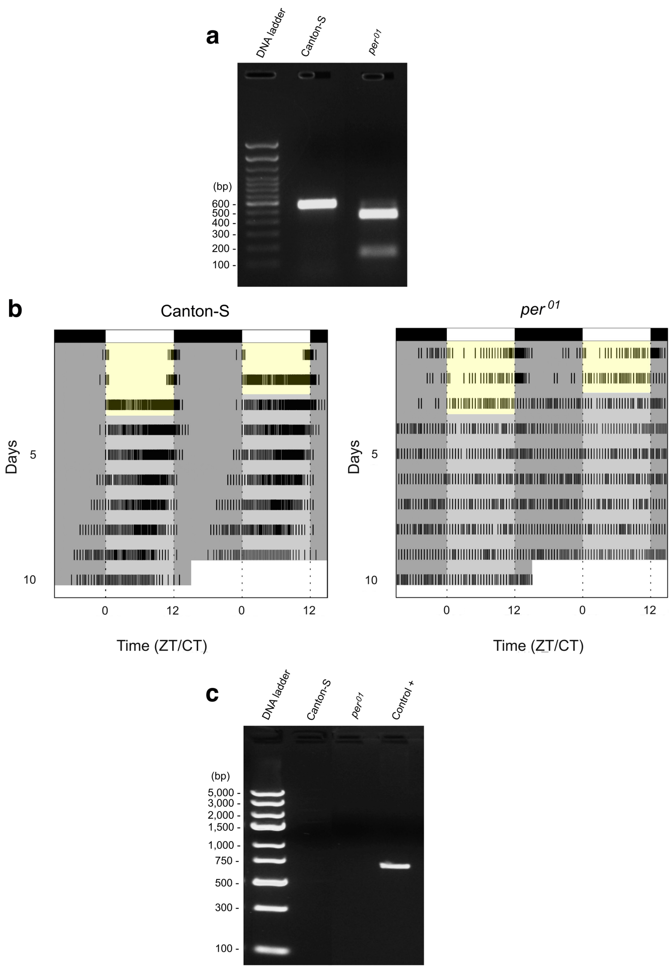


**Supplementary Figure S1**: **Genotyping, locomotor activity, and *Wolbachia* spp. screening in wild-type and *per*$^{01}$ flies.** (**a**) Genotyping of wild-type (Canton-S) and *per*$^{01}$ flies. A single 551 bp DNA fragment ("Canton-S") or two digestion products (125 and 426 bp) ("*per*$^{01}$") indicate the presence of *per*$^{+}$ and *per*$^{01}$ alleles in Canton-S and *per*$^{01}$ flies, respectively. (**b**) Double plotted actograms showing the locomotor activity of Canton-S (N = 87, left panel) and *per*$^{01}$ males (N = 27, right panel). After three days in 12:12 LD, flies were subjected to DD conditions for seven days. The light and dark phases are represented by yellow and grey areas, respectively. Periodogram analysis detected rhythmicity in 97.7% of Canton-S flies and 22.2% of *per*$^{01}$ mutants. (**c**) Bacterial endosymbiont *Wolbachia* PCR screening. In Canton-S and *per*$^{01}$ lanes*,* absence of PCR amplicons indicates that Canton-S and *per*$^{01}$ flies were *Wolbachia*-free. The band in "Control +" shows a ~ 600 bp *Wolbachia* amplification product obtained from gDNA of a known *Wolbachia*-infected fly line. In (**a**) and (**c**), DNA ladder show 100 bp and 1 kb DNA markers, respectively.

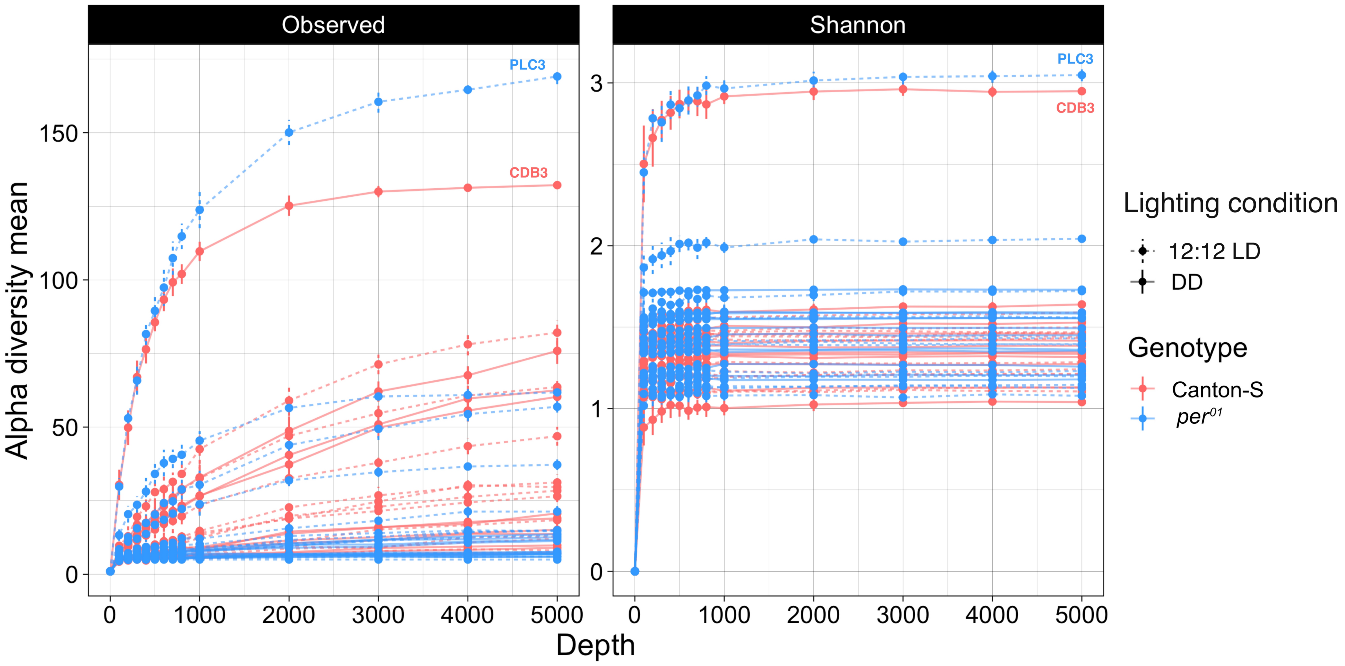


**Supplementary Figure S2**: **Rarefaction curves of samples based on alpha diversity measures.** Number of observed ASVs (left) and Shannon index (right) plotted *versus* the number of sequencing strips, randomly extracted from each sample. Data were normalized to the 90% of the smallest sample size (sample code PLB2). Canton-S and *per*$^{01}$ samples are shown in red and blue, respectively. Dashed and solid lines indicate 12:12 LD and DD conditions, respectively.

```
Lactiplantibacillus plantarum A     TAGGGAATCTTCCACAATGGACGAAAGTCTGATGGAGCAACGCCGCGTGAGTGAAGAAGG     60
Lactiplantibacillus plantarum B     TAGGGAATCTTCCACAATGGACGAAAGTCTGATGGAGCAACGCCGCGTGAGTGAAGAAGG     60
Lactiplantibacillus plantarum C     TAGGGAATCTTCCACAATGGACGAAAGTCTGATGGAGCAACGCCGCGTGAGTGAAGAAGG     60
                                    ************************************************************

Lactiplantibacillus plantarum A     GTTTCGGCTCGTAAAACTCTGTTGTTAAAGAAGAACATATCTGAGAGTAACTGTTCAGGT     120
Lactiplantibacillus plantarum B     GTTTCGGCTCGTAAAACTCTGTTGTTAAAGAAGAACATATCTGAGAGTAACTGTTCAGGT     120
Lactiplantibacillus plantarum C     GTTTCGGCTCGTAAAACTCTGTTGTTAAAGAAGAACATATCTGAGAGTAACTGTTCAGGT     120
                                    ************************************************************

Lactiplantibacillus plantarum A     ATTGACGGTATTTAACCAGAAAGCCACGGCTAACTACGTGCCAGCAGCCGCGGTAATACG     180
Lactiplantibacillus plantarum B     ATTGACGGTATTTAACCAGAAAGCCACGGCTAACTACGTGCCAGCAGCCGCGGTAATACG     180
Lactiplantibacillus plantarum C     ATTGACGGTATTTAACCAGAAAGCCACGGCTAACTACGTGCCAGCAGCCGCGGTAATACG     180
                                    ************************************************************

Lactiplantibacillus plantarum A     TAGGTGGCAAGCGTTGTCCGGATTTATTGGGCGTAAAGCGAGCGCAGGCGGTTTTTTAAG     240
Lactiplantibacillus plantarum B     TAGGTGGCAAGCGTTGTCCGGATTTATTGGGCGTAAAGCGAGCGCAGGCGGTTTTTTAAG     240
Lactiplantibacillus plantarum C     TAGGTGGCAAGCGTTGTCCGGATTTATTGGGCGTAAAGCGAGCGCAGGCGGTTTTTTAAG     240
                                    ************************************************************
                                                                            *
Lactiplantibacillus plantarum A     TCTGATGTGAAAGCCTTCGGCTCAACCGAAGAAGTGCATCGGAAACTGGGAAACTTGAGT     300
Lactiplantibacillus plantarum B     TCTGATGTGAAAGCCTTCGGCTCAACCGAAGAAGTGCATCGGAAACTGGGAAACTTGAGT     300
Lactiplantibacillus plantarum C     TCTGATGTGAAAGCCTTCGGCTCAACCGAAGAAGTGCATTGGAAACTGGGAAACTTGAGT     300
                                    *************************************** ********************
                                      *
Lactiplantibacillus plantarum A     GCAGAAGAGGACAGTGGAACTCCATGTGTAGCGGTGAAATGCGTAGATATATGGAAGAAC     360
Lactiplantibacillus plantarum B     GTAGAAGAGGACAGTGGAACTCCATGTGTAGCGGTGAAATGCGTAGATATATGGAAGAAC     360
Lactiplantibacillus plantarum C     GCAGAAGAGGACAGTGGAACTCCATGTGTAGCGGTGAAATGCGTAGATATATGGAAGAAC     360
                                    * **********************************************************

Lactiplantibacillus plantarum A     ACCAGTGGCGAAGGCGGCTGTCTGGTCTGTAACTGACGCTGAGGCTCGAAAGTATGGGTA     420
Lactiplantibacillus plantarum B     ACCAGTGGCGAAGGCGGCTGTCTGGTCTGTAACTGACGCTGAGGCTCGAAAGTATGGGTA     420
Lactiplantibacillus plantarum C     ACCAGTGGCGAAGGCGGCTGTCTGGTCTGTAACTGACGCTGAGGCTCGAAAGTATGGGTA     420
                                    ************************************************************

Lactiplantibacillus plantarum A     GCAAAC     426
Lactiplantibacillus plantarum B     GCAAAC     426
Lactiplantibacillus plantarum C     GCAAAC     426
                                    ******
```

**Supplementary Figure S3**: **Alignment of the three ASVs attributed to *Lactiplantibacillus plantarum*.** The three ASVs are indicated as *L. plantarum* A, B and C. Red asterisks show nucleotide variations at positions 280 (*L. plantarum* C) and 302 (*L. plantarum* B) in the 16S DNA sequence. The alignment was generated using CLUSTAL 0 (version 1.2.4).

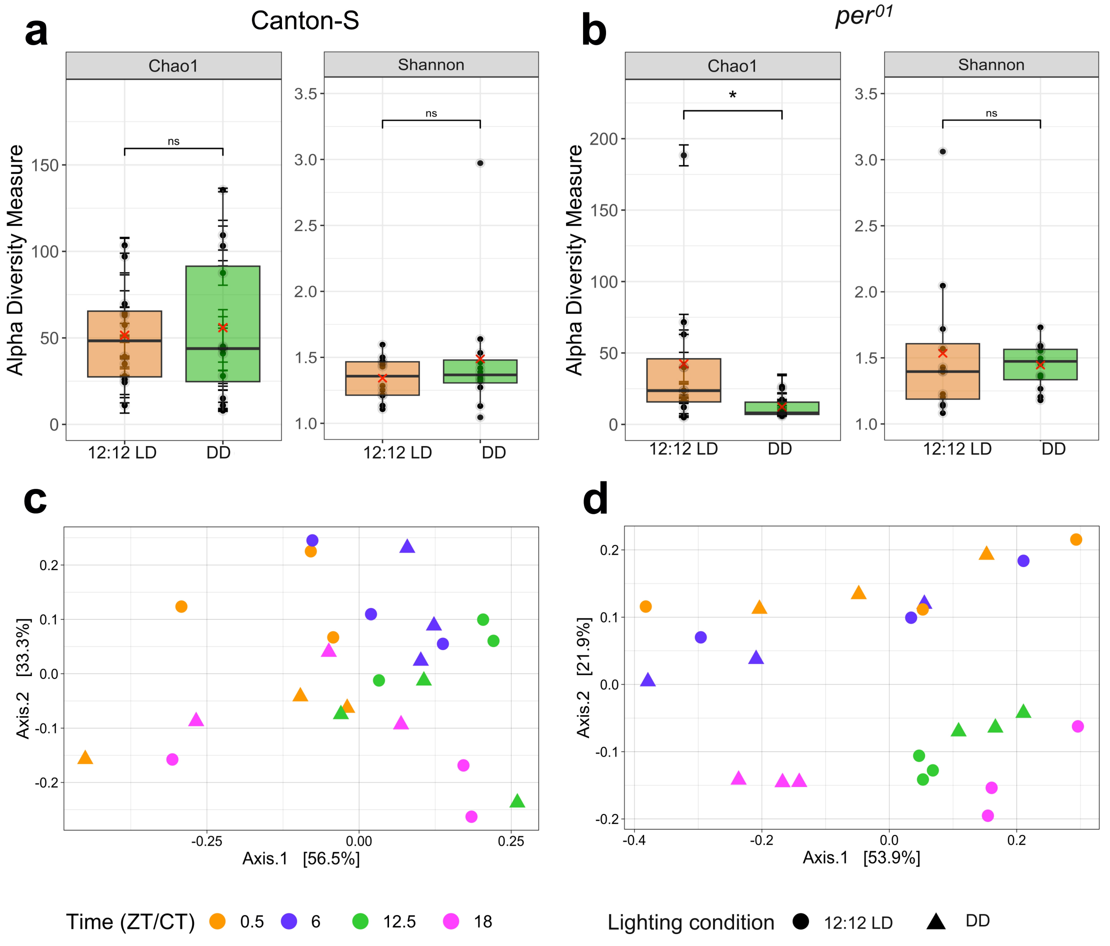


**Supplementary Figure S4**: **Effects of 12:12 LD and DD conditions on Canton-S and *per*$^{01}$ microbiota: alpha and beta diversity analyses.** (**a, b**) Shannon and Chao1 alpha diversity indices of Canton-S and *per*$^{01}$ microbiota under 12:12 LD and DD conditions. In each box plot, mean and median values are shown by red Xs, and horizontal black lines respectively; each point represents a biological replicate. In (**a**) Canton-S, no significant differences in Chao1 and Shannon indices were detected between 12:12 LD and DD conditions (Chao1: P = 0.908, ns; Shannon: P = 0.686, ns; Kruskal-Wallis test). In (**b**) *per*$^{01}$, Kruskal-Wallis test revealed significant differences between 12:12 LD and DD conditions in Chao1 index (P = 0.046) but not in Shannon index (P = 0.564, ns). (**c, d**) PCoA on Bray-Curtis matrix to measure the beta diversity between the 12:12 LD and DD regimes in both (**c**) Canton-S and (**d**) *per*$^{01}$ microbiota. For both host genotypes, PERMANOVA analyses did not show any significant dissimilarity in gut microbiota composition between 12:12 LD and DD conditions [(**c**) Canton-S: P = 0.444, ns; (**d**) *per*$^{01}$ P = 0.088, ns]. Colors indicate different time points: orange = ZT/CT 0.5, purple = ZT/CT 6, green = ZT/CT 12.5, pink = ZT/CT 18.

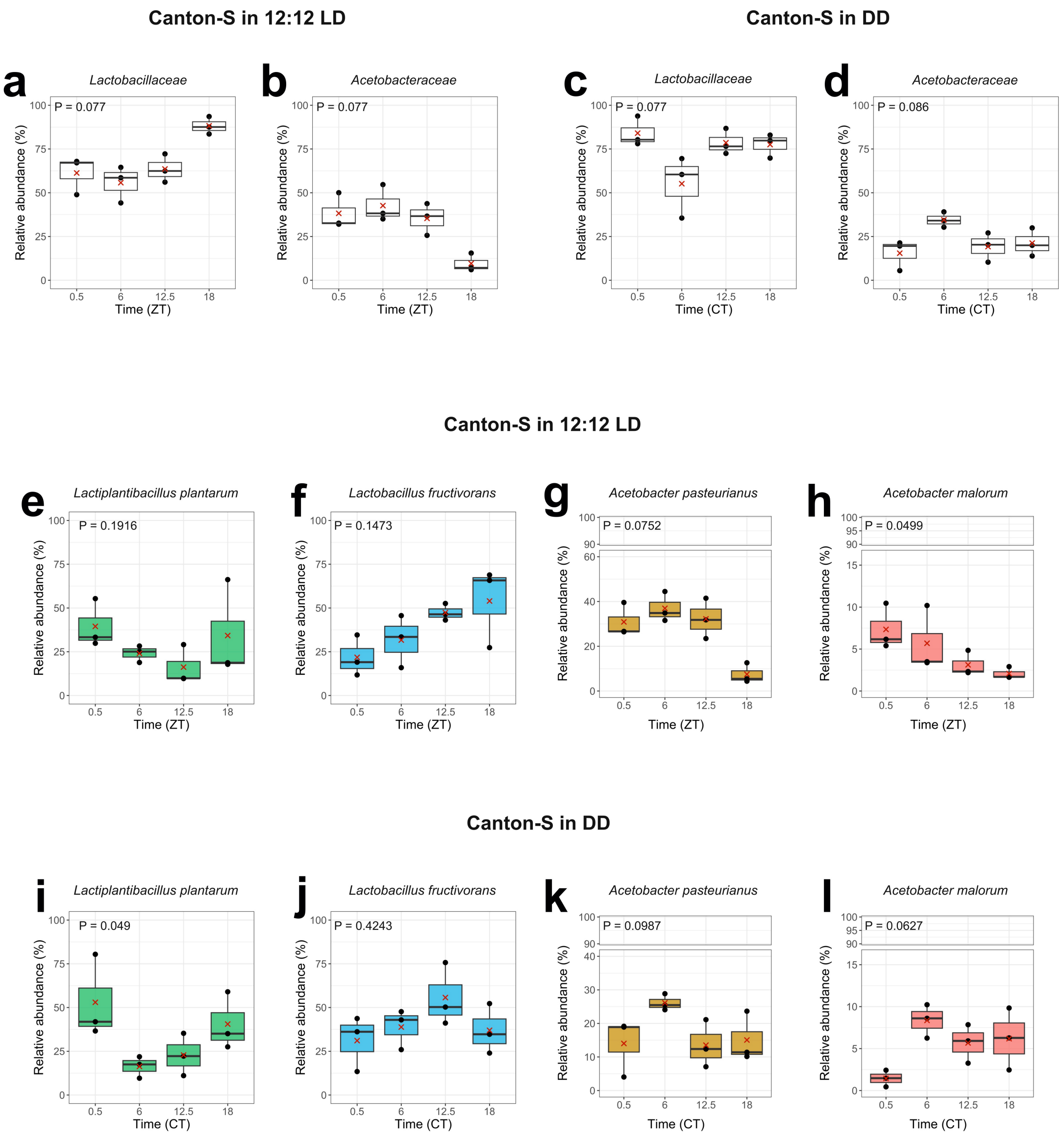


**Supplementary Figure S5: Daily relative abundances of the most prevalent bacterial families and ASVs in Canton-S microbiota under 12:12 LD and DD conditions.** Daily relative abundances (%) of *Lactobacillaceae* in (**a**) 12:12 LD, (**c**) DD, and *Acetobacteraceae* in (**b**) 12:12 LD and (**d**) DD conditions. (**e-l**) Daily relative abundances (%) of the most prevalent ASVs in (**e-h**) 12:12 LD, and (**i-l**) DD regimes. Mean and median values are indicated with a red cross and a solid black line, respectively. Kruskal-Wallis test revealed significant daily variations in (**h**) *A. malorum* in 12:12 LD conditions (P = 0.049) and in (**i**) *L. plantarum* in DD regimes (P = 0.049). All the other comparisons resulted P > 0.05, ns. P-values are reported in each panel.

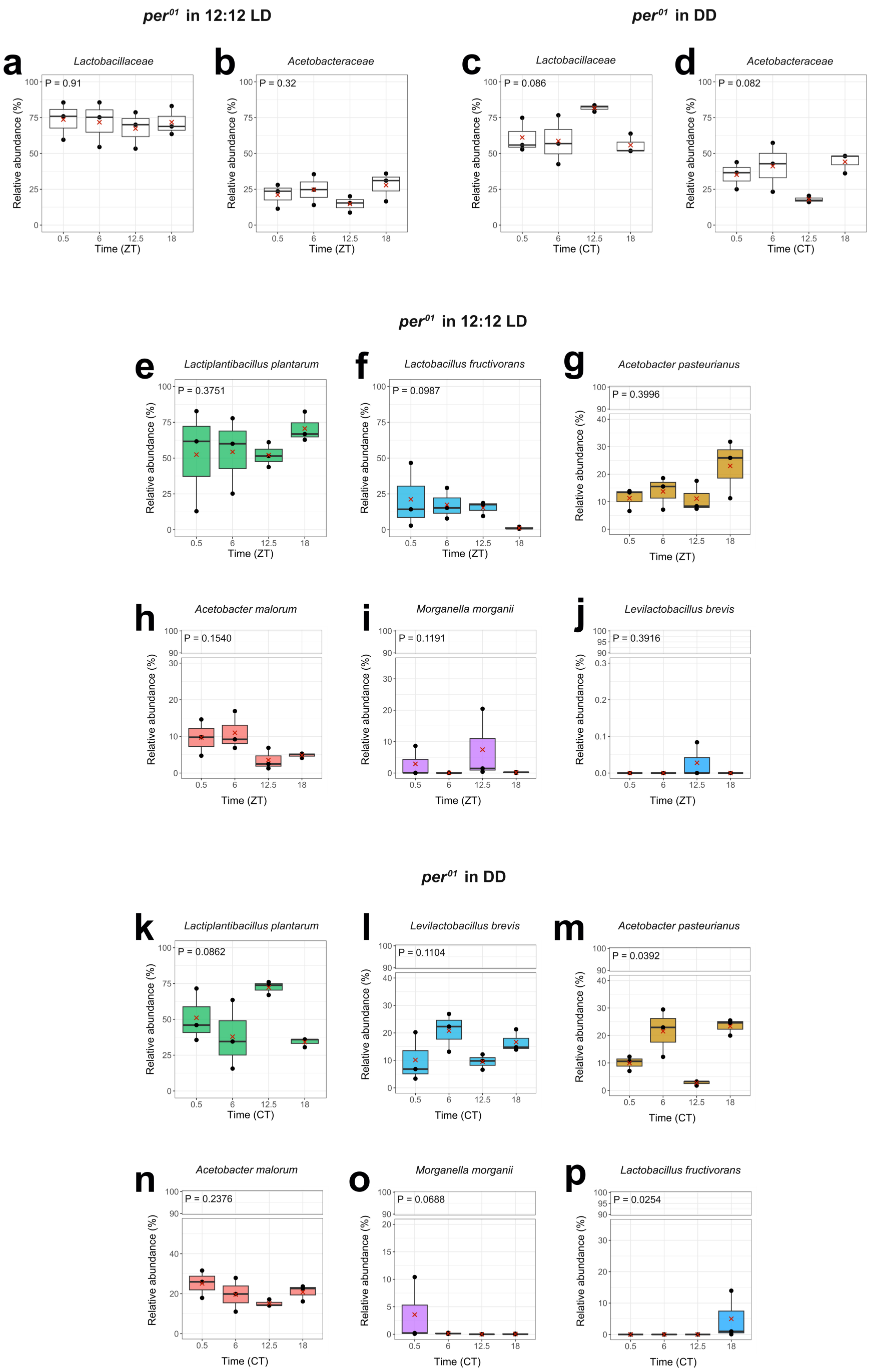


**Supplementary Figure S6: Daily relative abundances of the most prevalent bacterial families and ASVs in *per*$^{01}$ microbiota, under 12:12 LD and DD conditions.** Daily relative abundances (%) of *Lactobacillaceae* in (**a**) 12:12 LD, (**c**) DD, and *Acetobacteraceae* in (**b**) 12:12 LD and (**d**) DD conditions. (**e-p**) Daily relative abundances (%) of the most prevalent ASVs in (**e-j**) 12:12 LD, and (**k-p**) in DD regimes. Mean and median values are indicated with a red cross and a solid black line, respectively. Kruskal-Wallis test revealed significant daily variations in (**m**) *A. pasteurianus* (P = 0.039) and (**p**) *L. fructivorans* (P = 0.025), in DD regimes. All the other comparisons resulted P > 0.05, ns. P-values are reported in each panel.

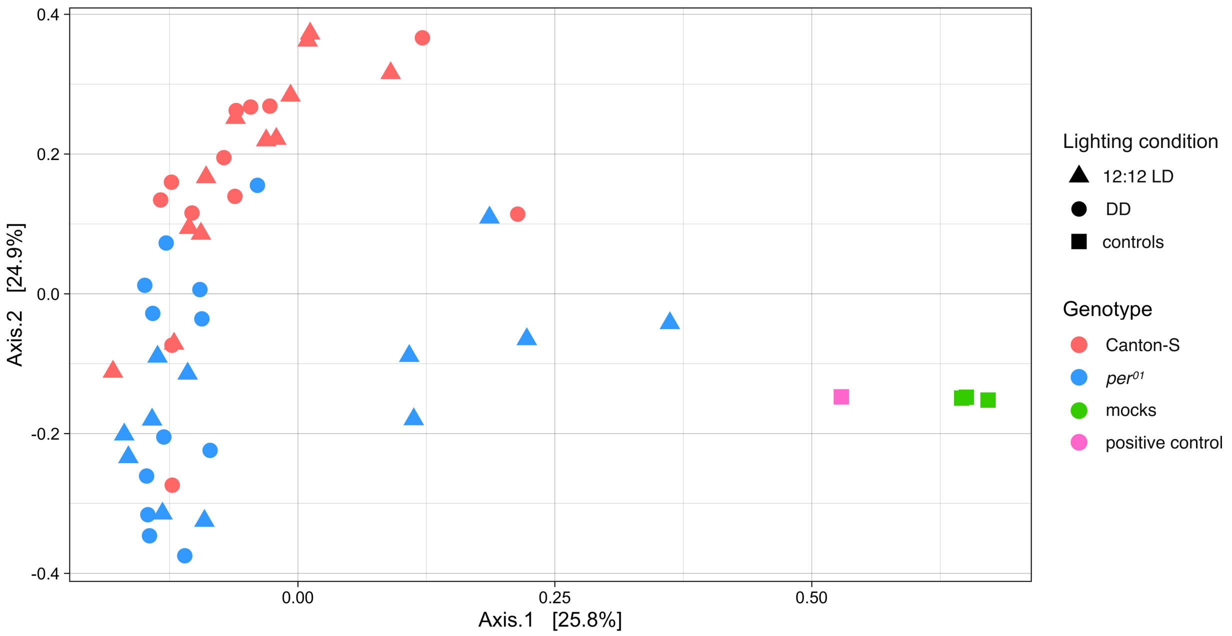


**Supplementary Figure S7: PCoA of Bray-Curtis distance matrix on raw data.** Canton-S and $per^{01}$ genotypes, as well as negative and positive controls are shown with different colors (red: Canton-S, blue: $per^{01}$, green: mocks, pink: positive control). 12:12 LD and DD conditions are indicated with different symbols (triangle =12:12 LD, circle = DD). The three negative controls (mocks) had a significantly different microbial composition compared to gut microbiota samples (PERMANOVA, P = 0.001).
